# A novel ciprofloxacin analogue enables daptomycin-mediated killing of resistant *Staphylococcus aureus* by increasing septum formation

**DOI:** 10.64898/2026.08.24.746612

**Authors:** Amber Y. Sefton, Elita Jauneikaite, Kam Pou Ha, Ravi Singh, Jacob Bradbury, Brigitte Lamy, Frédéric Laurent, Edward W. Tate, Thomas Lanyon-Hogg, Andrew M. Edwards

## Abstract

Quinolone antibiotics such as ciprofloxacin inhibit DNA gyrase, leading to DNA double-strand breaks that result in rapid bacterial killing and induction of the mutagenic SOS DNA repair response. By contrast, the ciprofloxacin analogue IMP-1700 inhibits ciprofloxacin-induced SOS, suggesting a novel mechanism of action. Here, we provide evidence that IMP-1700 targets the quinolone binding domain of DNA gyrase but triggers a significantly higher frequency of division septa in *S. aureus* compared with other DNA gyrase targeting antibiotics, including ciprofloxacin. In keeping with this finding, the lipopeptide antibiotic daptomycin, which targets the division septum, bound more strongly to IMP- 1700-treated cells relative to *S. aureus* exposed to other DNA gyrase inhibitors, leading to increased bacterial killing. This finding extended to a panel of paired daptomycin susceptible and resistant clinical isolates. We conclude that the ciprofloxacin analogue IMP-1700 has distinct effects on the cell envelope of *S. aureus*, despite appearing to share the same target as the parent drug, which result in the resensitisation of daptomycin resistant bacteria to the lipopeptide antibiotic.

## Introduction

*Staphylococcus aureus* is responsible for a wide range of different infections that affect people in both community and healthcare settings, including those of the skin, bloodstream, heart, bones and joints [1]. Treatment of staphylococcal infections is typically based on beta-lactamase resistant penicillins such as flucloxacillin [1,2]. However, resistant strains, known as methicillin-resistant *S. aureus* (MRSA) are globally disseminated and are now the second leading cause of drug-resistant bacterial infections worldwide [3].

Treatment of infections caused by MRSA relies on second-line antibiotics such as daptomycin or vancomycin, although resistance to each of these drugs can emerge during therapy via spontaneous mutations [4]. As such, there is a pressing need to develop new approaches to combat MRSA infections.

Fluoroquinolone antibiotics are broad-spectrum antibacterial agents and are amongst the most used classes used globally [5]. These drugs target the type II DNA topoisomerase enzyme complexes DNA gyrase and topoisomerase IV, which are crucial for bacterial control of DNA topology, DNA replication and cell division [5,6,7,8]. DNA gyrase is an A_2_B_2_ heterotetramer formed of two GyrA and two GyrB subunits and shares common architecture with the type IIA family of topoisomerases, including topoisomerase IV [9]. This enzyme is also a heterotetramer formed of ParE and ParC subunits that are homologous to gyrase GyrB and GyrA, respectively [6,9]. Comparably to GyrA, ParC is responsible for binding DNA and the enzyme’s catalytic properties, whereas, ParE acts in a similar manner to GyrB by controlling ATP binding [6,9]. Both DNA gyrase and topoisomerase IV use a strand passage mechanism which involves performing transient double stranded cleavage in one molecule of DNA, then passing a second DNA duplex through the double-strand break before religation [6,7,8,9,10]. Despite their similarity, DNA gyrase and topoisomerase IV perform different functions *in vivo*. Gyrase works in front of the replication fork and is responsible for the introduction of negative supercoils to compact the DNA, whereas topoisomerase IV performs ATP-dependent decatenation after replication to disentangle newly replicated DNA and allow daughter chromosome segregation [11,12].

Inhibition of DNA gyrase and topoisomerase IV by fluoroquinolone antibiotics blocks DNA replication and leads to DNA double strand breaks, which triggers the SOS DNA repair pathway [10,13]. Previous work developed a ciprofloxacin analogue, IMP-1700, which had potent antibacterial activity against methicillin susceptible *S. aureus* SH1000 and also potentiated ciprofloxacin activity, including against a quinolone resistant strain, USA300 JE2 [14] (Fig. 1A). Furthermore, IMP-1700 suppressed induction of the SOS DNA repair pathway by ciprofloxacin [14,15].

**Figure 1.**
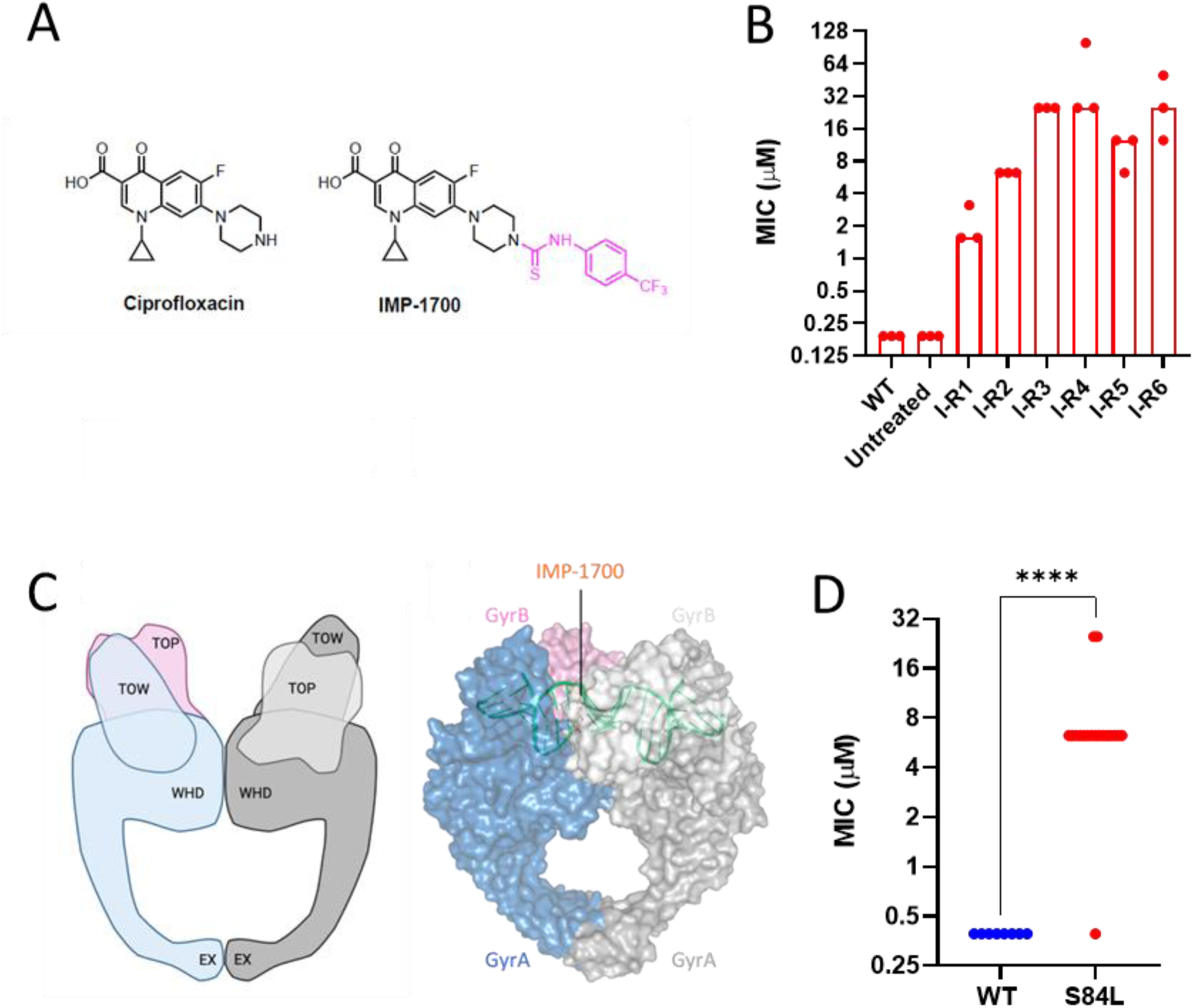
Selection for decreased IMP-1700 susceptibility results in mutations within *gyrA* that confer cross resistance to ciprofloxacin. **(A)** chemical structures of ciprofloxacin and IMP-1700. **(B)** susceptibility of independently generated *S. aureus* SH1000 isolates from the resistance-selection assay as determined by the MIC of IMP1700. Wild-type (WT) is the original strain and untreated refers to an isolate that was passaged in the absence of IMP-1700. I-R1 through I-R5 were isolated after passage with IMP-1700. N=3 independent experiments **(C)** diagram and space-filling model (PDB: 2XCT) of *S. aureus* DNA gyrase highlighting key domains and the proposed binding site of IMP-1700 based on the location of the S84L substitution that conferred decreased susceptibility. GyrB, pink; GyrA, blue; DNA, green; proposed IMP-1700 binding site indicated. **(D)** IMP-1700 MIC values for a collection of clinical isolates with wild-type (WT) DNA gyrase or strains containing the S84L substitution. N=3 independent experiments. Data analysed by Student’s t-test. **** p = <0.001.

In purified protein assays, analogues of IMP-1700 inhibit the nuclease activity of the AddAB nuclease/helicase complex that processes DNA double strand breaks for repair [16]. IMP-1700- functionalised Sepharose can pull down the staphylococcal AddAB DNA repair enzyme (known as RexAB in this organism) from bacterial lysate, supporting RexAB as the target of this compound [14]. These findings are consistent with work showing that loss of RexAB increases staphylococcal susceptibility to ciprofloxacin and suppresses the SOS response triggered by this antibiotic and others [13,14,17,18,19,20]. However, there is no direct evidence for engagement of RexAB by IMP-1700 in live cells, and since RexAB is not essential, the antibacterial activity of IMP-1700 must occur via another target [21].

Recent work has suggested that, in *E. coli* at least, the target of IMP-1700 could be DNA gyrase [22]. In support of this, IMP-1700 is less active against the JE2 strain of *S. aureus* which is ciprofloxacin resistant via a substitution in DNA gyrase, relative to the SH1000 strain which is susceptible to the fluoroquinolone. However, IMP-1700 did not show inhibitory activity in a purified recombinant *E. coli* DNA gyrase supercoiling assay [14]. Therefore, the aim of this work was to determine the mode of action of IMP-1700 on growth inhibition of *S. aureus*.

### Mutations in *gyr*A confer reduced susceptibility to IMP-1700

As a first step towards understanding how IMP-1700 inhibits staphylococcal growth, we selected for isolates with reduced susceptibility by serial passage in the presence of increasing concentrations of the compound (Supplementary Figure S1). We used the well characterised *S. aureus* SH1000 strain since this has a high degree of susceptibility to both IMP-1700 and ciprofloxacin [14,23] (Supplementary Table S1).

The minimum inhibitory concentration (MIC) of IMP-1700 was 0.2 µM for wild type *S. aureus* SH1000 and did not change during 8 serial passages in the absence of the compound (Fig. 1A,B). However, in the presence of IMP-1700, six isolates were recovered after 8 serial passages, with MICs that ranged from 1.56-25 µM (8-125-fold increase) (Fig. 1B).

Whole genome sequencing revealed that all isolates had mutations in the *gyrA* gene, resulting in a serine to leucine substitution at position 84 (S84L) of DNA gyrase subunit A, which is well established to confer ciprofloxacin resistance in *S. aureus* (Supplementary Table S2, Fig. 1C) [24,25]. Some isolates also had mutations in *parC* or *parE*, genes, which have also been linked to fluoroquinolone resistance [26] (Supplementary Table S2).

Isolate SH1000 I-R5 was selected for further analysis, since this isolate only differed from the wild type strain by a single SNP conferring the S84L substitution. Susceptibility testing of the SH1000 I -R5 isolate confirmed the expected increase in ciprofloxacin MIC, but there was no change in susceptibility to other DNA gyrase targeting drugs (zoliflodacin, novobiocin) or antibiotics that target cell wall production (fosfomycin, oxacillin), the membrane (daptomycin) or protein synthesis (tetracycline), underlining the specificity of this substitution for quinolone resistance (Supplementary Table S3).

Next, we tested whether the S84L substitution resulted in decreased IMP-1700 susceptibility in a panel of clinical isolates [27] (Supplementary Table S1). All 8 of the isolates examined that lacked the S84L substitution had IMP-1700 MICs of 0.4 µM, whereas 16/17 isolates with S84L had IMP-1700 MICs >4 µM (Fig. 1D). Further evidence for the importance of the S84L variant comes from the well characterised *S. aureus* strain JE2, which has this substitution and has been shown previously to have a higher IMP-1700 MIC than the SH1000 strain [14,28,29].

Taken together, these findings indicated that IMP-1700 and ciprofloxacin have the same or overlapping binding site in DNA gyrase (Fig. 1C), but they appear to have distinct effects on the enzyme in *S. aureus* [14].

### IMP-1700 does not inhibit staphylococcal RexAB/AddAB

Next, we investigated whether IMP-1700 inhibited RexAB, which has previously been suggested to be a target [14]. We firstly examined whether there was synergy between IMP-1700 and ciprofloxacin against wild-type *S. aureus* and a *rexB*::Tn mutant that lacks functional RexAB. This revealed synergy between IMP-1700 and ciprofloxacin in both wild-type and *rexB*::Tn mutants, indicating that synergy was not due to RexAB inhibition (Supplementary Fig. S2).

Since bacteria often have multiple DNA repair systems, which could conceivably compensate for the loss of RexAB in the mutant [13], we further tested the impact of IMP-1700 on RexAB by measuring helicase, nuclease and ATPase activity of recombinant staphylococcal RexAB as described previously [18]. However, we found no inhibition of any of the activities assayed. Taken together, these assays failed to provide evidence that RexAB is a target for IMP-1700 (Supplementary Fig. S2).

### IMP-1700 disrupts the staphylococcal cell cycle, leading to increased septation

Topoisomerase II enzymes such as DNA gyrase play a crucial role in the bacterial cell cycle during DNA replication and chromosome segregation, particularly by controlling DNA topology [30,31]. To further investigate DNA gyrase as a target of IMP-1700, we used microscopy to examine its impact on the cell cycle.

Work by Monteiro *et al.* (2015) characterized the cell cycle of *S. aureus*, which is sequential and occurs over three consecutive cycles of division (phases) in three orthogonal planes [32]. Phase 1 cells do not have septum formation as they have recently divided. Initiation of septum synthesis is observed in phase 2 cells. Cells in phase 3 exhibit a complete septum as the mother cell is about to split into two daughter cells, which re-enter phase 1 of the cell cycle once split [32].

Staining of cells with BODIPY-vancomycin enabled visualisation of the peptidoglycan cell wall, including the division septum, enabling determination of the cell cycle phase. In keeping with previous work, ∼60 % wild type SH1000 cells were in growth phase 1 in the absence of treatment, with ∼40% of cells in phases 2 or 3 (Fig. 2A,B) [32]. We then repeated the analysis with *S. aureus* treated with 2 X MIC of IMP-1700 (0.4 µM). This concentration was used to account for the larger number of cells present in these assays relative to those used to determine the MIC.

**Figure 2.**
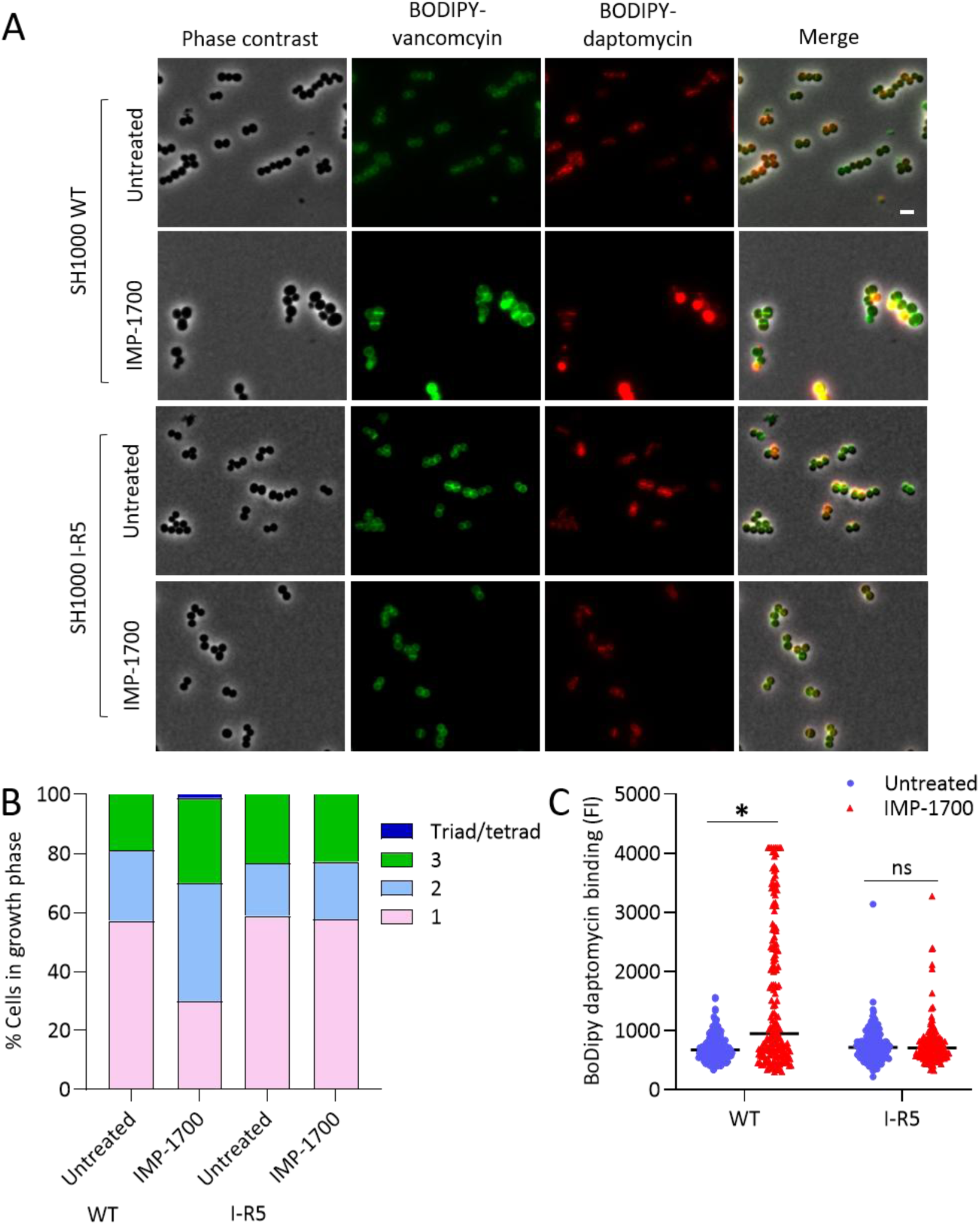
IMP-1700 causes increased septation in *S. aureus*. **(A)** *S. aureus* SH1000 WT and SH1000 IMP-1700-I-R5 strains were left untreated or exposed to 2 X the wild type MIC of IMP-1700 (0.4 µM) for 2 h before co-staining with BODIPY-daptomycin or BODIPY-vancomycin and analyses were performed by phase contrast and fluorescence microscopy; scale bar, 2 µm. **(B)** BODIPY-vancomycin- stained cells were used to quantify growth phase of 60 cells per replicate (180 cells total) and data are presented as the mean percentage of cells in each growth phase. Phase 1, no septum; phase 2, septum synthesis initiated; phase 3, complete septum formation; triad/tetrad, two or three complete septum per mother cell, respectively. **(C)** BODIPY-daptomycin binding of 60 cells per replicate (180 cells total) was quantified by fluorescence intensity and represented in the graph as a median and analysed by a two-way ANOVA with Sidak’s post hoc test (*P < 0.0001).

Cells exposed to IMP-1700 had markedly different levels of BODIPY-vancomycin staining, relative to untreated bacteria, and the number of cells in phase 1 decreased to ∼30 %, with concomitant increases in the frequency of cells in phases 2 and 3, as well as the appearance of cells with triad or tetrad septation (Fig. 2A,B). To understand if the effect of IMP-1700 on the staphylococcal cell involved DNA gyrase, we repeated the experiment with the SH1000 IMP-1700 I-R5 isolate using the same concentrations of inhibitor as for the wild type. In the absence of IMP-1700, this isolate had a similar cell cycle profile to the wild type, with ∼ 60 % cells in growth phase 1, but crucially, this did not change in the presence of IMP-1700 at the same concentration as that used for experiments with the wild type (Fig. 2A,B).

Daptomycin is a lipopeptide antibiotic that targets phosphatidylglycerol in the staphylococcal membrane, primarily at the division septum, leading to disrupted cell wall synthesis [33,34]. Given the increased septation caused by IMP-1700, we hypothesised this would lead to increased binding of daptomycin and tested this by attaching a BODIPY label to the antibiotic as described previously [35]. As hypothesised, IMP-1700 promoted binding of daptomycin to SH1000 wild type cells relative to untreated controls (Fig. 2A,C). However, IMP-1700 had no effect on daptomycin binding to the SH1000 IMP-1700 I-R5 isolate when used at the same concentration as for the wild type (Fig. 2A,C).

Taken together, these findings showed that IMP-1700 caused increased septation in wild-type *S. aureus* SH1000 but not the I-R5 isolate, indicative of a role for DNA gyrase.

### IMP-1700-mediated increased septation and daptomycin binding is not a common property of DNA gyrase inhibitors

Although the S84L substitution reduced the susceptibility of *S. aureus* to IMP-1700, it was still possible to inhibit bacterial growth using a higher concentration of the inhibitor (Supplementary table S3) [14]. This is important because MRSA strains, for which new treatments are needed, often have the S84L substitution in DNA gyrase [24,25]. Therefore, we next tested the impact of IMP-1700-mediated growth inhibition on the MRSA strain JE2, again using 2 X MIC of IMP-1700 (10 µM) before bacteria were stained with BODIPY-vancomycin to visualise the cell wall, and Hoescht dye to visualise the nucleoid as an additional marker of septation.

In keeping with previous work, and similar to *S. aureus* SH1000, 62 % of untreated *S. aureus* JE2 cells were in growth phase 1 and contained a single chromosome, with 38 % in phases 2 and 3 (Fig. 3 A,B,C). In contrast, <20 % of *S. aureus* JE2 cells exposed to IMP-1700 were in growth phase 1. We also observed ∼10 % of cells existing as tetrads, which was not seen for untreated cells (Fig. 3 A,B).

**Figure 3.**
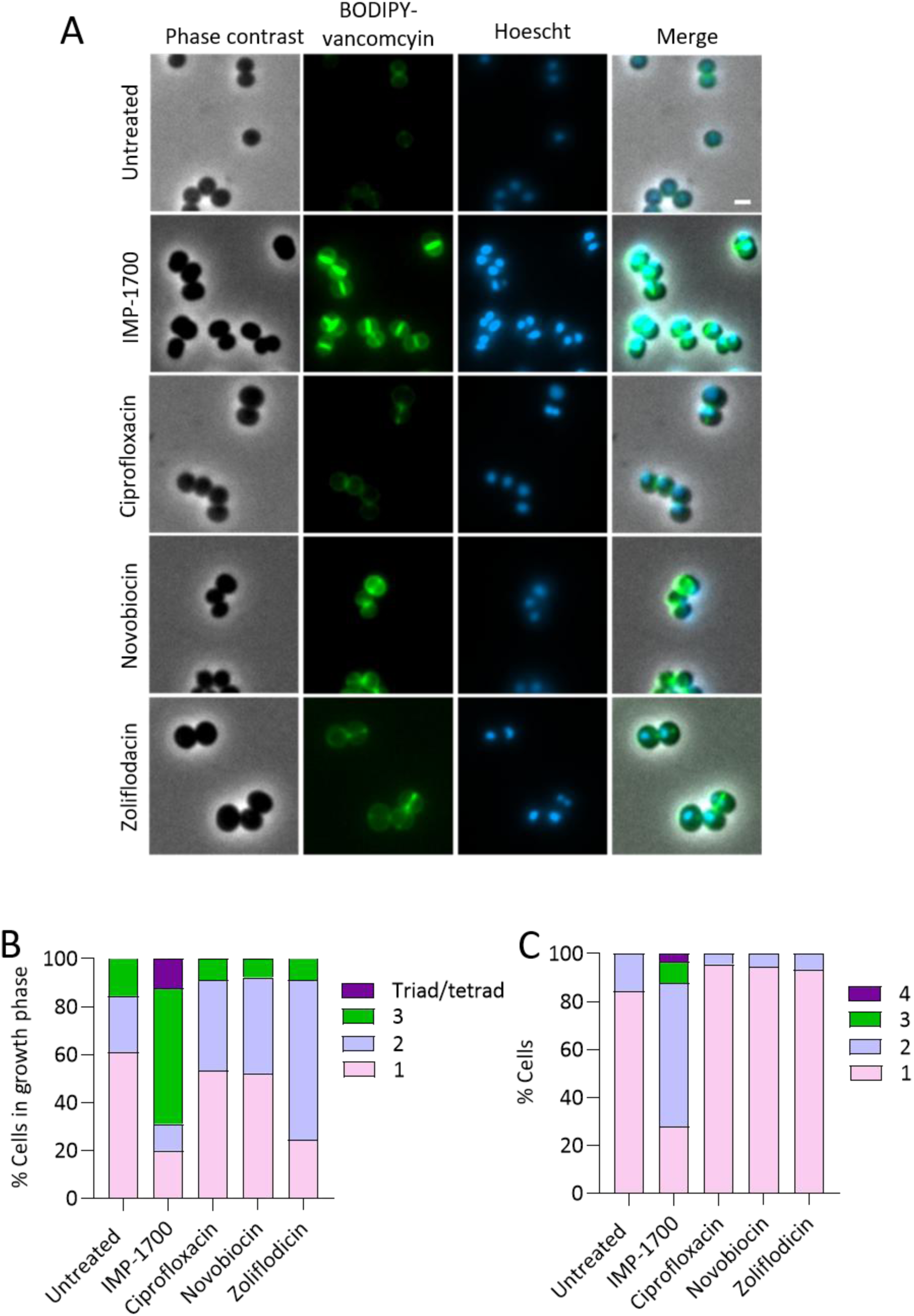
IMP-1700 causes increased septation and abnormal nucleoid formation in MRSA JE2. **(A)** Phase contrast and fluorescence microscopy of *S. aureus* JE2 WT cells left untreated or exposed to 2x MIC IMP-1700, ciprofloxacin, novobiocin or zoliflodacin for 2 h before co-staining with BODIPY-vancomycin and Hoechst; scale bar, 1 µm. **(B)** Growth phase of BODIPY-vancomycin stained cells quantified: phase 1, no septum; phase 2, septum synthesis initiated; phase 3, complete septum formation; triad/tetrad, two or three complete septum per mother cell, respectively. **(C)** Hoechst staining used to quantify number of chromosomes per mother cell. Data in **(B,C)** are represented as the mean percentage of 30 cells per replicate (90 total).

Next, we sought to determine whether IMP-1700 triggered increases in septation were a common feature of DNA gyrase inhibitors by comparison with cells treated with 2 X MIC ciprofloxacin, novobiocin or zoliflodacin [36,37]. In contrast to IMP-1700, treatment of *S. aureus* with ciprofloxacin or novobiocin had very little effect on the distribution of growth phases relative to untreated cells, whereas zolifloidacin exposure resulted in a larger proportion of cells in growth phase 2 (> 60 %) relative to I (24 %) or III (8 %) (Fig. 3 A,B).

The chromosome morphology of IMP-1700 treated cells was fully segregated, with no unsegregated chromosomes bisected with septa (Fig. 3 A). Interestingly, the cells with triad and tetrad septum formation also had 3 or 4 fully segregated chromosomes per mother cell, respectively (Fig. 3 A,C), indicating that new rounds of DNA replication and chromosome segregation occur in IMP-1700-treated cells. This result was unexpected as impairment of the topoisomerase II enzymes has been shown to regulate chromosome segregation and cell cycle progression, as observed here with ciprofloxacin, novobiocin and zoliflodacin (Fig. 3 A,B,C) [38,39].

As an additional confirmation of increased septum formation in bacteria exposed to IMP-1700, the fluorescent D-alanine analogue HADA was used to track peptidoglycan synthesis [40]. This showed significantly increased HADA incorporation at the septum of IMP-1700 cells relative to untreated cells (Supplementary Fig. S3). This was most likely due to synthesis, rather than a lack of hydrolysis, since oxacillin reduced HADA incorporation at the septum (Supplementary Fig. S3). We then confirmed increased daptomycin binding to IMP-1700 treated cells of *S. aureus* JE2 relative to untreated bacteria or those exposed to ciprofloxacin (Supplementary Fig. S4).

Taken together, these data demonstrate that IMP-1700 causes changes in growth phase, septum formation, nucleoid distribution and daptomycin binding in an MRSA strain with the S84L substitution, relative to untreated cells. These effects were not seen with other types of DNA gyrase targeting antibiotics. Therefore, despite the structural similarity of IMP-1700 to ciprofloxacin, these two compounds cause very different effects on the *S. aureus* cell cycle.

### IMP-1700 enhanced daptomycin binding occurs primarily at the division septum

To better understand whether increased binding of daptomycin to IMP-1700 treated cells was due to increased septation, we undertook detailed localisation of daptomycin binding with reference to the division septum. As expected from the work described above, the septa of IMP-1700 treated wild type *S. aureus* JE2 cells bound significantly more BODIPY-vancomycin than untreated cells or those incubated with ciprofloxacin, indicative of more peptidoglycan (Fig. 4A). IMP-1700-treated cells also exhibited high levels of BODIPY-daptomycin binding at the septum than untreated or ciprofloxacin treated cells (Fig. 4B). However, there was also enhanced BODIPY-daptomycin binding at the cell periphery of IMP-1700 treated cells compared to untreated or ciprofloxacin treated bacteria, indicative of dispersal of the antibiotic from the septum [33] (Fig. 4B). Combined, these findings confirmed that the increased binding of daptomycin to IMP-1700 treated *S. aureus* is primarily associated with the increased degree of septation that occurred in these cells, but not those treated with ciprofloxacin.

**Figure 4.**
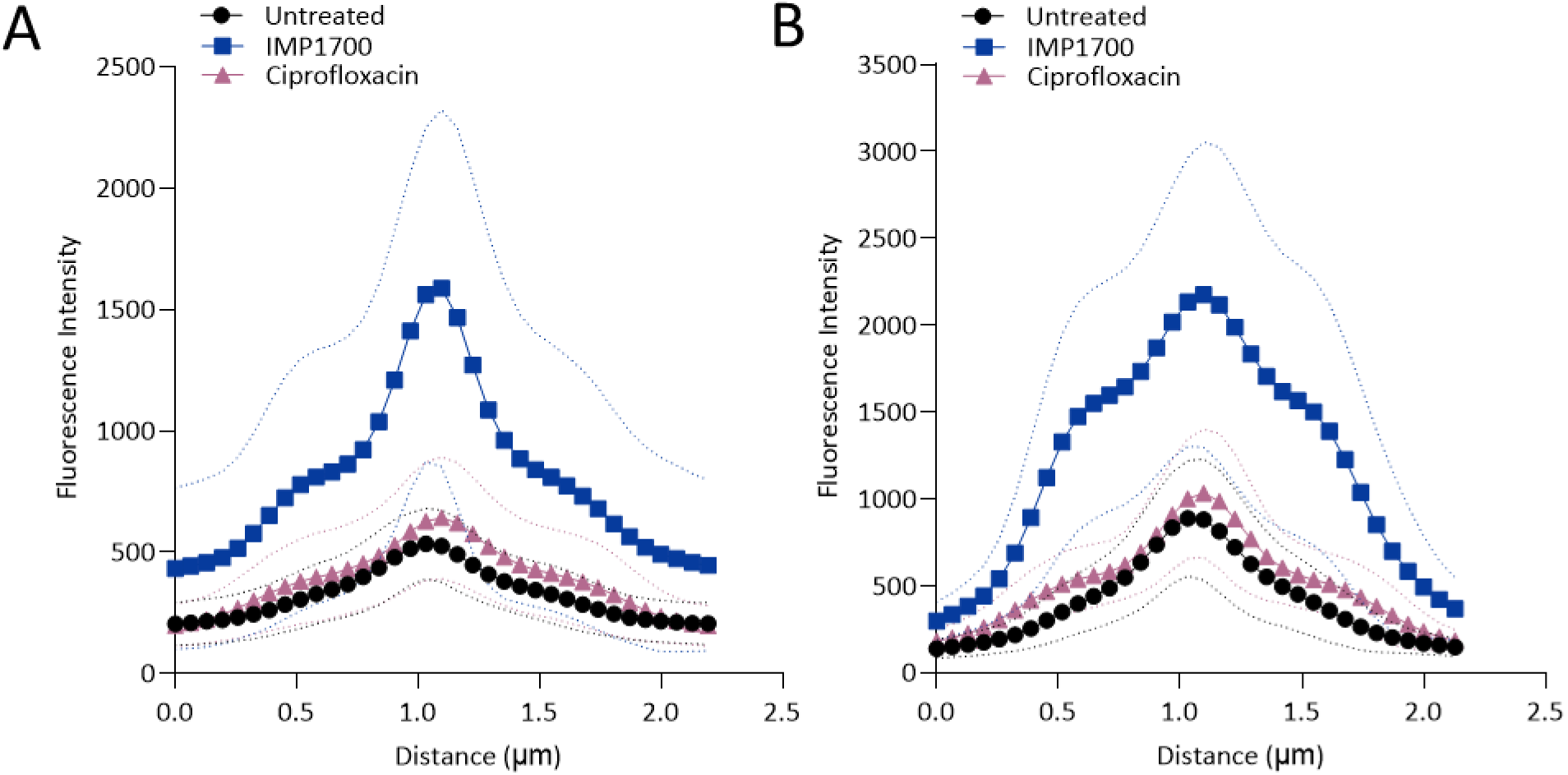
Enhanced daptomycin binding correlates with increased septation in IMP-1700-treated cells. (A,B) Profiling of individual *S. aureus* JE2 WT cells for peptidoglycan abundance (BODIPY-vancomycin, **A**) and daptomycin binding (BODIPY-daptomycin, **B**) to cells. Graphs show the fluorescence intensity of a cross-section perpendicular to the septum of cells stained with **(A)** BODIPY-vancomycin or **(B)** BODIPY-daptomycin. Data in both panels represent the mean of 20 cells per replicate (60 cells total, N=3). Standard deviations of the mean are shown by dotted line rather than error bars to enhance clarity. Data were analysed by two-way ANOVA. Values for IMP-1700 treated cells were greater than those of untreated or ciprofloxacin treated bacteria at every point (p = <0.05).

### IMP-1700 promotes daptomycin-mediated killing of clinical daptomycin non-susceptible strains

Staphylococcal infections are often hard to treat, even in the case of drug susceptible strains, resulting in a growing interest in combination therapies to improve patient outcomes [1,2]. Therefore, we examined whether the IMP-1700-mediated increase in daptomycin binding translated into increased bacterial killing.

We firstly exposed *S. aureus* SH1000 wild type to a range of concentrations of IMP-1700, or ciprofloxacin as a comparator, in the absence of daptomycin and measured bacterial viability via CFU counts after 8 hours. Both IMP-1700 and ciprofloxacin caused a dose-dependent reduction in viability, with ∼1000-fold reduction in CFU counts when used at the highest concentration of 12.5 µM (Fig. 5A).

**Figure 5.**
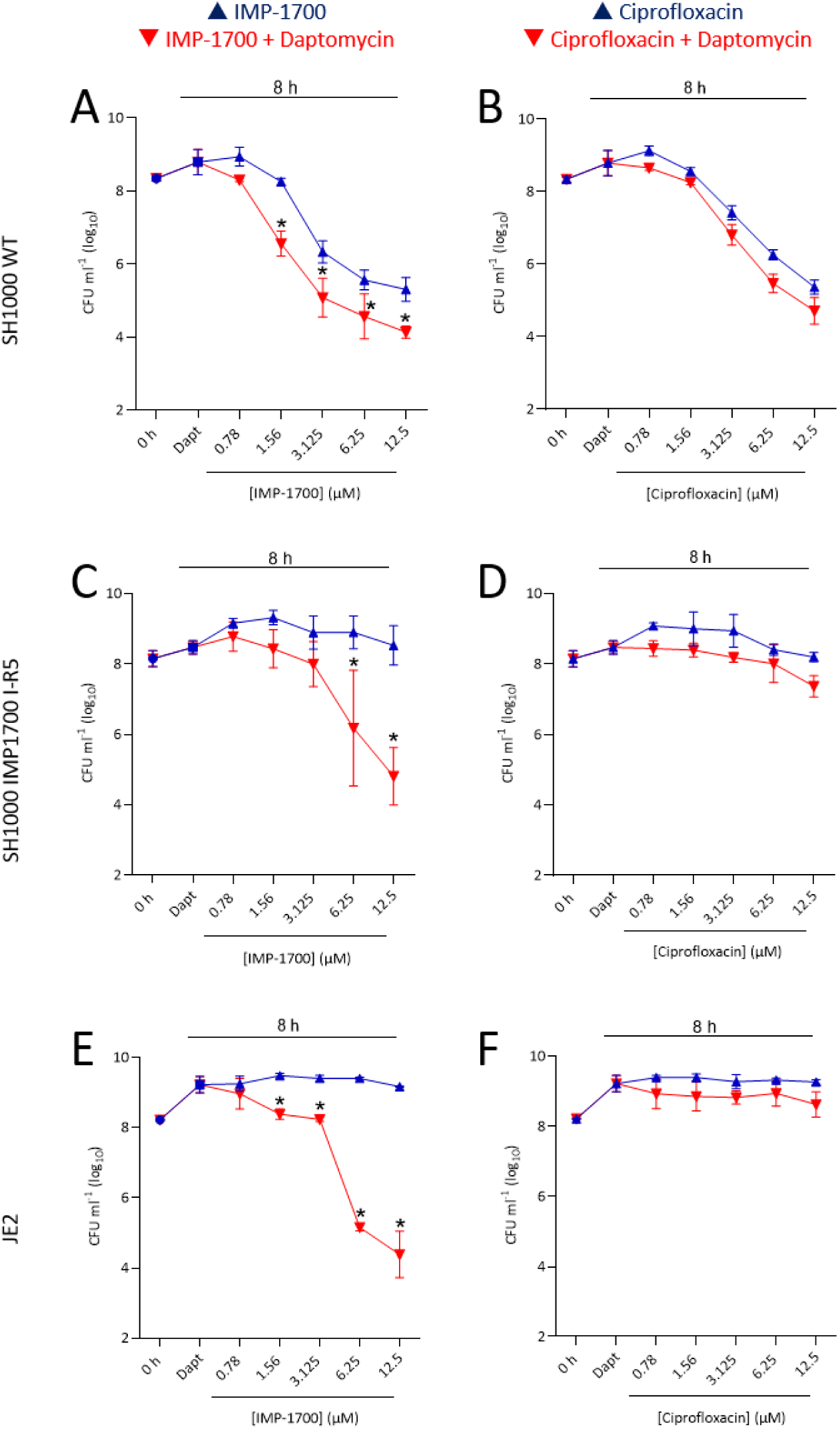
IMP-1700 potentiates daptomycin killing of *S. aureus*. Survival of SH1000 WT **(A,B)**, SH1000 I-R5 **(C,D)** or JE2 WT **(E,F)** exposed to a range of either IMP-1700 **(A,C,E)** or ciprofloxacin **(B,D,F)** across a range of concentrations either with or without 5 µg ml^-1^ daptomycin after 8 h. 0 h = CFU counts at the start of the assay; Dapt = daptomycin alone after 8 h of treatment. Data are presented as the mean (n=3 *±*standard deviation) and analysed by two-way ANOVAs with Tukey’s post-hoc test (*P < 0.05; comparison of compound alone vs compound with daptomycin).

At the concentration used (5 µg ml^-1^), daptomycin alone did not reduce CFU counts. However, CFU counts were significantly lower when IMP-1700 (1.56 µM and higher concentrations) was combined with daptomycin, compared to IMP-1700 alone (Fig. 5A). By contrast, CFU counts were not significantly different between ciprofloxacin alone versus ciprofloxacin combined with daptomycin at any of the concentrations examined (Fig. 5B). Therefore, IMP-1700 showed synergistic bactericidal activity with daptomycin, whilst ciprofloxacin did not.

Next, we repeated this experiment with the *S. aureus* SH1000 I-R5 isolate with decreased IMP-1700 susceptibility. At 1.56 µM and 3.13 µM, IMP-1700 failed to synergise with daptomycin, in contrast to *S. aureus* SH1000 wild type (Fig. 5A,C). However, there was significant promotion of daptomycin-mediated killing at the higher IMP-1700 concentrations of 6.25 and 12.5 µM, whereas ciprofloxacin had no effect on daptomycin-mediated bacterial killing at any concentration tested (Fig. 5C,D). Since daptomycin is typically used to treat infections caused by MRSA strains that often have the S84L substitution in DNA gyrase [27,28], this finding suggests that it may be possible to use IMP-1700 to enhance daptomycin activity in these strains, albeit at a higher dose than for strains without the substitution.

To explore whether IMP-1700 could potentiate daptomycin activity against MRSA with the S84L substitution in DNA gyrase, we repeated the IMP-1700-daptomycin synergy assay with *S. aureus* JE2. As for SH1000 I-R5, IMP-1700 alone did not inhibit the growth of *S. aureus* JE2, but there was synergistic bactericidal activity with daptomycin, which was similar in magnitude to that seen with wild type SH1000 at higher concentrations, whilst ciprofloxacin had no effect (Fig. 5E,F).

To understand whether this finding with *S. aureus* JE2 extended to other MRSA strains with the DNA gyrase S84L substitution we examined a panel of clinical isolates that consisted of matched pairs of daptomycin-susceptible and daptomycin non-susceptible isolates. For all 15 isolates, IMP-1700 showed synergistic bactericidal activity with daptomycin, demonstrating a consistent effect on MRSA strains, even when they are non-susceptible to daptomycin (Supplementary Fig. S5). In addition, using MIC checkerboard assays with IMP-1700 and daptomycin, we determined the fractional inhibitory concentration index (FICI) and found synergistic growth inhibitory activity for 10 of 16 isolates with the DNA gyrase S84L substitution and additive inhibitory effects in the other 6 isolates (Supplementary Table S4).

Combined, these data show that IMP-1700 alone lacks bactericidal activity against strains with the DNA gyrase S84L substitution, but maintains synergy with daptomycin against these isolates, even in the case of daptomycin-non-susceptible strains.

## Discussion

Previous work showed that IMP-1700 had growth inhibitory activity against *S. aureus* and suppressed the SOS DNA repair pathway triggered by ciprofloxacin [14]. However, the mechanism by which IMP-1700 inhibits staphylococcal growth was unknown.

By selecting for decreased susceptibility to the growth-inhibitory activity of IMP-1700 across parallel cultures, we found evidence that DNA gyrase is targeted. The most consistent mutations selected for cause a S84L substitution that is commonly found in clinical isolates with resistance to ciprofloxacin [24,25]. In keeping with this, analysis of a large panel of clinical isolates found significantly decreased IMP-1700 susceptibility in isolates with this substitution, relative to isolates with wild type DNA gyrase. These findings agree with a previous study that showed reduced IMP-1700 susceptibility in an *E. coli* isolate with *gyrA* and *parC* mutations [22].

Given the structural similarities between IMP-1700 and ciprofloxacin, it is not surprising that DNA gyrase was found to be a target. What is surprising, however, is the very different phenotypic consequences of ciprofloxacin and IMP-1700, including divergent SOS responses and impacts on cellular replication, leading to increased septation and susceptibility to the lethal activity of the antibiotic daptomycin when used in combination with IMP-1700. This divergence may be due to subtle differences in the way IMP-1700 and ciprofloxacin bind DNA gyrase, or due to the presence of a secondary target. Recent work has described IMP-1700 analogues with enhanced growth-inhibitory and SOS-suppressing activity [15]. One of these, OXF-077, has since been shown to target staphylococcal signal peptise IB (SpsB) that enables protein trafficking to the membrane and is required for induction of SOS [41]. Earlier work showed that loss of signal peptidase function leads to cell cycle arrest and aberrant cell separation, resulting in a significant increase in cells in growth phase 3 and associated increase in septation, similar to what we observed with IMP-1700 [42]. Therefore, it is possible that IMP-1700 also acts via signal peptidase or another secondary target besides DNA gyrase. However, it is also possible that the divergent effects of ciprofloxacin and IMP-1700 arise via subtle differences in the way they bind DNA gyrase, which will require detailed structural analyses to elucidate. For example, previous work indicated that nalidixic acid, the first synthetic quinolone antibiotic, caused increased septation in *S. aureus*, which indicates that relatively minor differences in quinolone structure can cause distinct effects on replication [43].

As daptomycin is one of very few treatment options for MRSA infection, significant fundamental and clinical research has recently focused on combining daptomycin with other antibiotics to improve treatment outcomes [4,44,45,46]. Much of this work has involved beta-lactams and other drugs that inhibit peptidoglycan synthesis and thereby synergise with daptomycin’s impact on cell wall synthesis activity [44,45,46,47,48,49]. Furthermore, inhibition of peptidoglycan synthesis prevents cell wall accumulation under *in vivo* conditions, which reduces daptomycin activity [50].

Unfortunately, however, despite promising data from in *vitro* and animal experiments, as well as increased infection clearance rates in patients, there has been a lack of improvement in patient survival from antibiotic combination therapy, suggesting that more work is needed to develop effective approaches [45,46,49,51]. Whilst it is far too early to propose a combination therapy based on daptomycin and IMP-1700, the observed potentiation of antibiotic activity of ciprofloxacin [12] and daptomycin, as well as inhibition of the SOS response induced by oxacillin and trimethoprim [41], indicates that the IMP-1700 series may hold significant potential as an antibiotic adjuvant, which warrants further detailed mechanistic investigation of these potentiation mechanisms.

## Materials and Methods

### Bacterial strains and growth conditions

The bacterial strains used in this study are shown in Table 2.1. All strains listed were grown to stationary phase in 3 ml tryptic soy broth (TSB; BD Biosciences) in a 30 ml universal tube with shaking at 180 rpm at 37 °C or statically on tryptic soy agar plates (TSA; BD Biosciences) at 37 °C for ∼16 h. When required for selection, antibiotic was added to the growth media as stated. Strains were stored at -80 °C in a 1:1 ratio of overnight culture and 50% glycerol in cryovials.

### Determination of Minimum Inhibitory Concentrations

The MIC of an antibiotic or compound was determined using the broth microdilution assay as previously described [52]. This was performed in 96-well plates with the antibiotic/compound serially diluted two-fold in 95 µl TSB. Inoculation was made up to ∼1 x 10^5^ colony forming units (CFU) ml^-1^ by diluting stationary phase *S. aureus* 1:1000 and inoculating 5 µl into each well containing 95 µl media, resulting in a final volume of 100 µl. After 17 h static incubation at 37 °C, the MIC was determined by eye and defined as the lowest concentration of antibiotic or compound with no visible growth of bacteria. Assays to determine daptomycin MIC were performed in TSB containing 1.25 mM CaCl_2_.

### Antibiotic checkerboard assay

Checkerboard assays were used to determine whether IMP-1700 was synergistic, additive or indifferent [53] with a range of antibiotics tested against *S. aureus*. The broth microdilution method was used to set up two 96-well plates with 95 µl TSB containing two-fold serial dilutions of antibiotic or compound. One plate had two-fold dilutions of one compound/antibiotic running vertically and the second plate had the other compound/antibiotic running horizontally. It was important that the dilutions started at double the highest concentration to be tested and no antibiotic/compound was diluted into the final row/column. A second two-fold serial dilution was performed. This involved pipetting 95 µl from the last column of one plate containing the lowest concentration and mixing this with the identical column in the second plate before pipetting the doubly diluted media back to the initial plate. This was repeated across the whole plate until the column containing the highest concentration was doubly diluted. One of the plates containing 95 µl media per well was inoculated with a final inoculum of ∼1 x 10^5^ CFU ml^-1^ (5 µl of 1:1000 overnight culture). The inoculated plate was statically incubated at 37 °C for 17 h. A Bio-Rad iMark microplate reader (Bio-Rad Laboratories) was then used to read OD_595_ values. An OD_595_ reading < 0.1 was used to define no bacterial growth and the MIC was determined as the lowest concentration of antibiotic/compound to achieve this. The European Committee for Antimicrobial Susceptibility Testing (EUCAST) guidelines used a fractional inhibitory concentration (FIC) index to calculate whether there was synergy between two drugs [53].

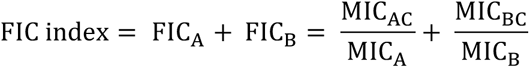

MIC_A_ = MIC of A alone; MIC_B_ = MIC of B alone; MIC_AC_ = MIC of A in combination; MIC_BC_ = MIC of B in combination. The EUCAST guidelines defined an FIC index ≤ 0.5 as synergy, ≤ 1 as additive and > 1 as indifferent. Daptomycin checkerboard assays were performed with 1.25 mM CaCl_2_ in the media.

### Selection for mutants with reduced IMP-1700 susceptibility via serial passage

The serial passage protocol was modified from methods described previously [54,55]. Eight independent colonies of *S. aureus* SH1000 were each grown in 3 ml TSB (180 rpm, 37 °C) for ∼16 h to stationary phase. Next, 100 µl of one of the overnight cultures was used to inoculate 3 ml TSB without IMP-1700 (untreated) and this culture was termed ‘SH1000 WT passaged’. Then 100 µl from each of the remaining seven cultures (subsequently named SH1000 intermediate, R1, R2, R3, R4, R5 and R6) was used to inoculate 3 ml TSB containing 0.5 x or 1 x MIC of IMP-1700. This resulted in a total of 14 cultures exposed to IMP-1700 and one culture left untreated, which were all incubated at 37 °C, 180 rpm for 24 h. The following day, the culture growing at the highest IMP-1700 concentration was used to make a glycerol stock and then for the subsequent passage grown in the same and also a two-fold higher concentration of IMP-1700 than the previous passage. This process was repeated for eight consecutive days of serial passaging with increasing IMP-1700 concentrations; the ‘SH1000 intermediate strain’ was passaged with IMP-1700 for three days; and the ‘SH1000 WT passaged’ strain was left untreated throughout the eight days of passaging as a control. The IMP-1700 MIC of cultures re-streaked from the glycerol stocks and grown in the absence of IMP-1700 were determined to confirm that resistance was stable.

### Whole-genome sequencing and genomic analysis

Isolates with decreased IMP-1700 susceptibility had DNA extracted using GenElute Bacterial Genomic DNA kit (Sigma-Aldrich) and sequenced at the Imperial BRC Genomics Facility. Raw sequencing files were assessed for quality using FastQC v0.11.6 (https://github.com/s-andrews/fastqc) and low quality reads were removed using Trimmomatic v0.39 (https://github.com/usadellab/trimmomatic) [56]. Bacterial species were verified using Kraken v2.1.2 (https://github.com/DerrickWood/kraken2) [57]. Snippy v4.6.0 (<u>github.com/tseemann/snippy</u>) was used to identify genetic variation compared to a reference genome NZ_CP020619.1 and the susceptible isolate SH1000 WT. Bioinformatics analyses were done using High Performance Computing at Imperial College London and using European Galaxy server (https://usegalaxy.eu). Illumina reads of SH1000 WT, R1-R6 isolates were deposited in European Nucleotide Archive (https://www.ebi.ac.uk/ena/browser/home) under BioProject PRJEB75336.

### Time-kill assays

This assay is based on a previous protocol [49]. An overnight culture of stationary phase *S. aureus* was washed x3 in PBS by alternate rounds of centrifugation at 17,000 x g for 2 min and resuspension. Washed *S. aureus* cells were diluted 1:10 in TSB containing 1.25 mM CaCl_2_ by using 300 µl to inoculate a total of 3 ml TSB in a universal tube containing the relevant antibiotic or compound, which gave a final inoculum of 3 X 10^8^ CFU ml^-1^. These cultures were incubated for 8 h (37 °C, 180 rpm). At 0, 2, 4 and 8 h, or at timepoints stated, 100 µl aliquots were taken and ten-fold serial dilutions were made in 200 µl PBS in 96-well plates. Next, 10 µl of each dilution was plated onto TSA by spotting and spreading with a sterile loop, before the agar plates were incubated for ∼16 h and colony forming units (CFUs) were counted. Only dilutions containing 10-100 colonies were counted.

### Fluorescent labelling of daptomycin with BODIPY

Daptomycin was labelled with one of these two fluorophores as described previously [58]: green BODIPY FL N-Hydroxysuccinimide (NHS) ester (Life Technologies) or red BODIPY 558/568 NHS ester (Thermo Fisher). This was performed by mixing 50 µl of 50 mg ml^-1^ daptomycin with 100 µl 10 mg ml^-1^ BODIPY in sodium bicarbonate buffer (pH 8.5, 0.2 M) to make a total volume of 1 ml. This mixture was incubated in the dark for 4 h at 37 °C. Unconjugated BODIPY was removed by dialysis against water using a Float-A-Lyser G2 device (Spectrum Labs; molecular weight cut-off at 0.1-0.5 kDa). Regular distilled water (∼400 ml) changes were made throughout dialysis, which was performed in the dark at 4°C for 24 h.

### Phase contrast and fluorescence microscopy

For staining with HADA, *S. aureus* cells were grown to stationary phase, washed and diluted to a final density of ∼10^8^ CFU ml^-1^ in 1 ml TSB supplemented with 100 µM HADA and either left untreated or containing 2x MIC of oxacillin or IMP-1700. Cultures were incubated for 2 h at 37 °C with shaking at 180 rpm before a 200 µl aliquot was taken and washed 4 times in PBS before resuspending in 4% paraformaldehyde (PFA) to fix cells, which were then kept in the dark at room temperature until they were analysed by microscopy.

For BODIPY-vancomycin or BODIPY-daptomycin staining, stationary phase *S. aureus* cells were washed and diluted 1:10 in 1.5 ml TSB to a final density of ∼10^8^ CFU ml^-1^. The media was either left untreated or supplemented with 2x MIC IMP-1700, ciprofloxacin, novobiocin or zoliflodacin. Cultures were incubated (37 °C, 180 rpm) for 2 h before 200 µl aliquots were taken and bacteria co-stained with the following: 1.25 mM CaCl_2_; 40 µg ml^-1^ red BODIPY-daptomycin and 2 µg ml^-1^ green BODIPY-vancomycin (Thermo Fisher). After 10 min incubation at room temperature in the dark, cells were washed 4 times in PBS by rounds of centrifugation (17, 000 x *g*, 2 min) and resuspension, before resuspension of the final pellet in 4% PFA. These were kept at room temperature in the dark until microscopy was performed.

For BODIPY-vancomycin and Hoechst staining, stationary phase *S. aureus* cells were washed and diluted 1:10 in 1.5 ml TSB either left untreated or supplemented with 2x MIC IMP-1700, ciprofloxacin, novobiocin or zoliflodacin and incubated for 2 h at 37 °C, 180rpm. Aliquots (200 µl) were taken and mixed with 2 µg ml^-1^ BODIPY-vancomycin and 2 µg ml^-1^ Hoechst for 10 min in the dark at room temperature. Next, cells were washed x4 in PBS and resuspended in 4% PFA before being stored in the dark at room temperature until analysis by microscopy.

### Quantification of fluorescence by microscopy

Microscope slides were prepared by dissolving agarose in water to 1.2% using a microwave and allowing the solution to cool before pipetting 500 µl onto a microscope slide and covering with parafilm to set. Once solidified, the parafilm was removed and 2 µl aliquots of bacterial suspensions were taken of fixed cells and spotted onto the agarose before placing a cover slip on top. Samples were kept in the dark until analysis by phase contrast and fluorescence microscopy images were taken using the Zeiss Axio Imager A1 microscope coupled to an AxioCam MRm and a 100x objective. The Zen 2012 software (blue edition) was used to process these images. Detection of the different dyes was achieved by the following: DAPI filter set for HADA and Hoechst; Texas red filter set for red BODIPY-daptomycin; GFP filter set for BODIPY-vancomycin. Zen 2012 software (blue edition) was used to quantify the fluorescence intensity of cells and FIJI software was used to quantify the fluorescence intensity profile of a line drawn perpendicular to the septum, which was plotted vs distance.

## Acknowledgements

All authors acknowledge the provision of strains by the Network on Antimicrobial Resistance in *Staphylococcus aureus* (NARSA) Program under NIAID/NIH contract no. HHSN272200700055C. The funders had no role in the study design, interpretation of the findings, or the writing of the manuscript.

Angelika Grundling (Imperial College London) is acknowledged for providing strains and useful conversations.

