## Supplementary data file for "A novel ciprofloxacin analogue enables daptomycin-mediated killing of resistant *Staphylococcus aureus* by increasing septum formation"

Contains supplementary tables S1-S4 and supplementary figures S1-S5.

| **Strain** | | **Description** | **Source** |
|---|---|---|---|
| SH1000 | | *rsbU+* derivative of 8325-4 | 22 |
| JE2 | | LAC strain derived from USA300 CA-MRSA, cured of | 28 |
| IMP-1700 | SH1000 untreated | SH1000 passaged untreated for 8 days | This study |
| challenged | SH1000 I-intermediate | SH1000 passaged with IMP-1700 for 3 days | This study |
| | SH1000 I-R2 | SH1000 passaged with IMP-1700 for 8 days | This study |
| | SH1000 I-R3 | SH1000 passaged with IMP-1700 8 days | This study |
| | SH1000 I-R4 | SH1000 passaged with IMP-1700 8 days | This study |
| | SH1000 I-R5 | SH1000 passaged with IMP-1700 8 days | This study |
| | SH1000 I-R6 | SH1000 passaged with IMP-1700 8 days | This study |
| ST20201745 | | Pair 1; daptomycin sensitive; SCCmecA +ve | 26 |
| ST20201688 | | Pair 1; daptomycin resistant; SCCmecA +ve | 26 |
| ST20171642 | | Pair 2; daptomycin sensitive; SCCmecA +ve | 26 |
| ST20171643 | | Pair 2; daptomycin resistant; SCCmecA +ve | 26 |
| ST20171644 | | Pair 2; daptomycin resistant; SCCmecA +ve | 26 |
| ST20141980 | | Pair 3; daptomycin sensitive; SCCmecA +ve | 26 |
| ST20141981 | | Pair 3; daptomycin resistant; SCCmecA +ve | 26 |
| ST20200611 | | Pair 4; daptomycin sensitive; SCCmecA -ve | 26 |
| ST20200408 | | Pair 4; daptomycin resistant; SCCmecA -ve | 26 |
| ST20200244 | | Pair 5; daptomycin sensitive; SCCmecA +ve | 26 |
| ST20200225 | | Pair 5; daptomycin resistant; SCCmecA +ve | 26 |
| ST20182004 | | Pair 6; daptomycin sensitive; SCCmecA -ve | 26 |
| ST20181963 | | Pair 6; daptomycin resistant; SCCmecA -ve | 26 |
| ST20150287 | | Pair 7; daptomycin sensitive; SCCmecA -ve | 26 |
| ST20150288 | | Pair 7; daptomycin resistant; SCCmecA -ve | 26 |
| ST20171810 | | Pair 9; daptomycin sensitive; SCCmecA +ve | 26 |
| ST20171811 | | Pair 9; daptomycin resistant; SCCmecA +ve | 26 |
| ST20160260 | | Pair 10; daptomycin sensitive; SCCmecA +ve | 26 |
| ST20160261 | | Pair 10; daptomycin resistant; SCCmecA +ve | 26 |
| ST20161115 | | Pair 11; daptomycin sensitive; SCCmecA +ve | 26 |
| ST20161098 | | Pair 11; daptomycin resistant; SCCmecA +ve | 26 |
| ST20200655 | | Pair 12; daptomycin sensitive; SCCmecA +ve | 26 |
| ST20200654 | | Pair 12; daptomycin resistant; SCCmecA +ve | 26 |
| ST20171659 | | Pair 13; daptomycin sensitive; SCCmecA +ve | 26 |
| ST20171658 | | Pair 13; daptomycin resistant; SCCmecA +ve | 26 |

**Supplementary Table S1.** Bacterial strains used in this study.

| CHROM | POS | Type | Effect | Gene | Product |
|---|---|---|---|---|---|
| **WT** | | | | | |
| | | - | - | - | - |
| **Untreated** | | | | | |
| | | **-** | **-** | **-** | **-** |
| **I-R1** | | | | | |
| NODE_2 | 252845 | SNP | Missense variant; c.251C>T; p.Ser84Leu | *gyrA* | GyrA |
| NODE_3 | 275332 | MNP | n/a | n/a | n/a |
| **I-R2** | | | | | |
| NODE_1 | 422294 | ins | Conservative inframe insertion; c.101_102insACG p.Lys34_Arg35insArg | *parE* | ParE |
| NODE_2 | 252845 | SNP | Missense variant; c.251C>T p.Ser84Leu | *gyrA* | GyrA |
| NODE_3 | 275332 | MNP | n/a | n/a | n/a |
| **I-R3** | | | | | |
| NODE_1 | 418766 | ins | Conservative inframe insertion; c.1638_1639insGAT p.Gln546_Asp547insAsp | *parC* | ParC |
| NODE_2 | 252845 | SNP | Missense variant; c.251C>T p.Ser84Leu | *gyrA* | GyrA |
| **I-R4** | | | | | |
| NODE_1 | 420169 | SNP | Missense variant; c.239C>A p.Ser80Tyr | *parC* | ParC |
| NODE_2 | 252845 | SNP | Missense variant; c.251C>T p.Ser84Leu | *gyrA* | GyrA |
| **I-R5** | | | | | |
| NODE_2 | 252845 | SNP | Missense variant; c.251C>T p.Ser84Leu | *gyrA* | GyrA |
| NODE_3 | 275332 | MNP | n/a | n/a | n/a |
| **I-R6** | | | | | |
| NODE_2 | 252845 | SNP | Missense variant; c.251C>T p.Ser84Leu | *gyrA* | GyrA |
| NODE_3 | 275332 | MNP | n/a | n/a | n/a |

**Supplementary Table S2.** Mutations and inferred substitutions identified by WGS of *S. aureus* isolates with reduced susceptibility to IMP-1700, relative to wild type SH1000.

CHROM – Chromosome. POS – position.

| Antibiotic | SH1000 WT | SH1000 I-R5 | Fold increase in I-R5 |
|---|---|---|---|
| IMP-1700 | 0.2 µM | 25 µM | 125 |
| Ciprofloxacin | 1.56 µM | 50 µM | 32 |
| Zoliflodicin | 0.125 µg ml$^{-1}$ | 0.125 µg ml$^{-1}$ | - |
| Novobiocin | 0.125 µg ml$^{-1}$ | 0.125 µg ml$^{-1}$ | - |
| Fosfomycin | 32 µg ml$^{-1}$ | 32 µg ml$^{-1}$ | - |
| Oxacillin | 0.125 µg ml$^{-1}$ | 0.125 µg ml$^{-1}$ | - |
| Daptomycin | 1 µg ml$^{-1}$ | 1 µg ml$^{-1}$ | - |
| Tetracycline | 0.25 µg ml$^{-1}$ | 0.25 µg ml$^{-1}$ | - |

**Supplementary Table S3. MIC values of a range of antibiotics for *S. aureus* SH1000 WT and I-R5** (n=3; data presented as the median).

| Strain | FICI | Interpretation |
|---|---|---|
| $1642^{S}$ | 0.25 | Synergy |
| $1643^{R}$ | 0.75 | Additive |
| $1980^{S}$ | 0.5 | Synergy |
| $1981^{R}$ | 0.36 | Synergy |
| $244^{S}$ | 0.5 | Synergy |
| $255^{R}$ | 0.36 | Synergy |
| $1810^{S}$ | 0.63 | Additive |
| $1811^{R}$ | 0.38 | Synergy |
| $260^{S}$ | 0.16 | Synergy |
| $261^{R}$ | 0.75 | Additive |
| $115^{S}$ | 0.19 | Synergy |
| $98^{R}$ | 0.5 | Synergy |
| $655^{S}$ | 0.5 | Synergy |
| $654^{R}$ | 0.75 | Additive |
| $1659^{S}$ | 0.75 | Additive |
| $1658^{R}$ | 0.75 | Additive |

**Supplementary Table S4. IMP-1700 synergises with daptomycin against most clinical *S. aureus* isolates.** The tables shows FICI values for IMP-1700 and daptomycin derived from checkerboard MIC assays.

FICI defines synergy as $\leq$ 0.5, additive as > 0.5-1, indifference as 1-4 and antagonism as > 4 [52].

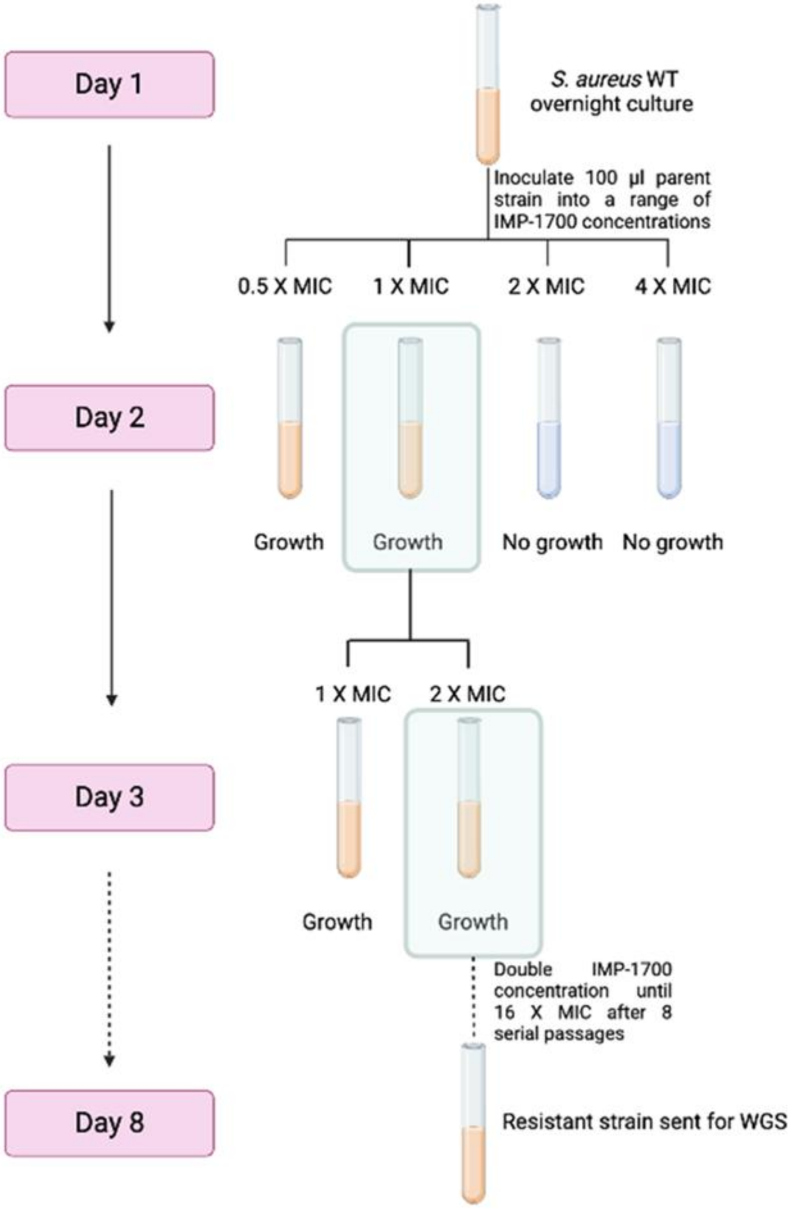

**Supplementary Figure S1. Selection for isolates with reduced IMP-1700 susceptibility.** Schematic showing an overview of the protocol used to passage *S. aureus* with IMP-1700 for eight days, resulting in the isolation of bacteria with reduced susceptibility. **Created with BioRender.com.**

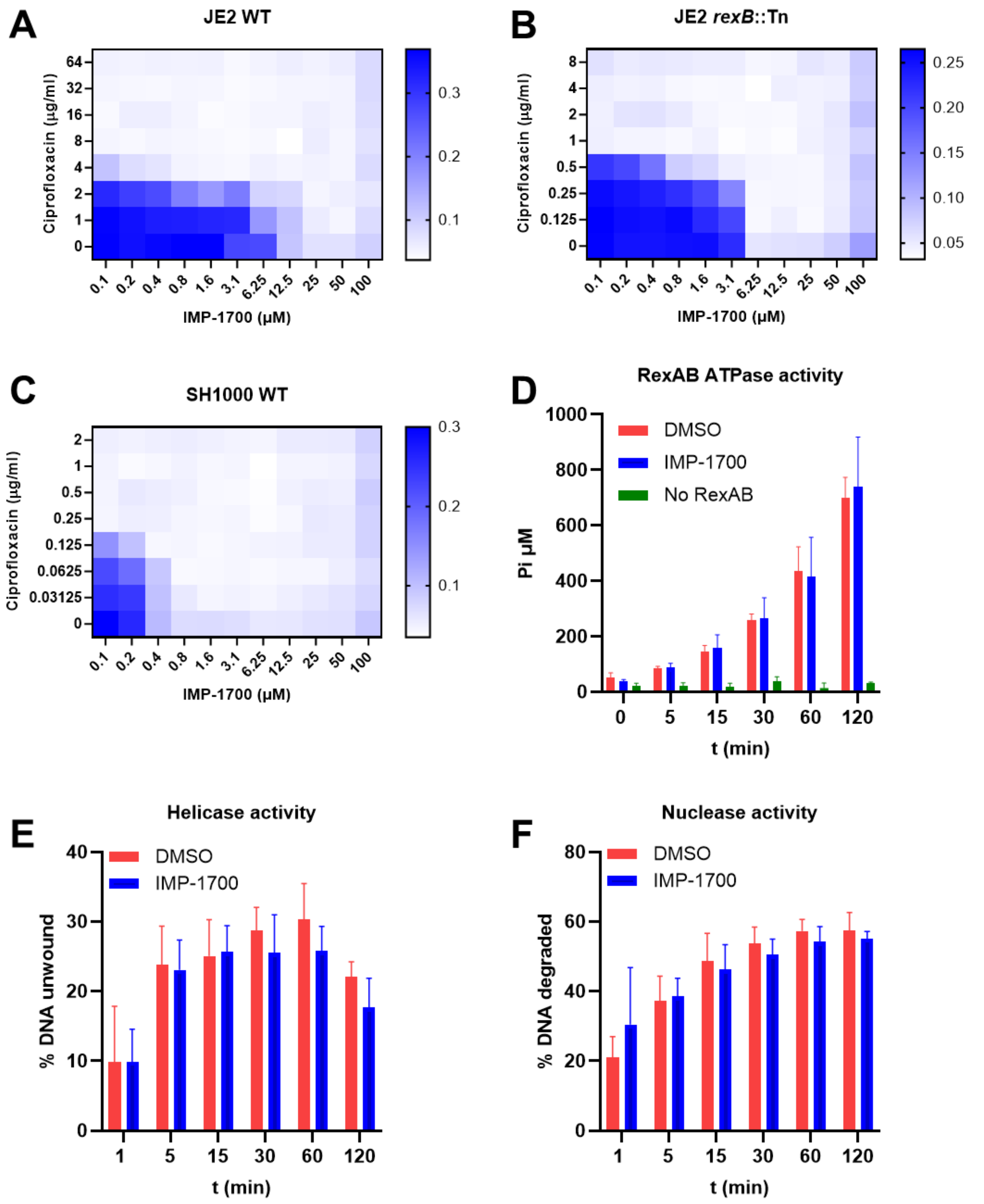

**Supplementary Figure S2. IMP-1700 does not inhibit RexAB.** Checkerboard assays to determine synergistic antibacterial activity of IMP-1700 and ciprofloxacin against *S. aureus* JE2 wild type (**A**), *S. aureus* JE2 *rexB*::Tn (**B**) or *S. aureus* SH1000 wild type (**C**). Recombinant RexAB ATPase (**D**), Helicase (**E**) or Nuclease (**F**) activity in the absence or presence of IMP-1700 (10 µM).

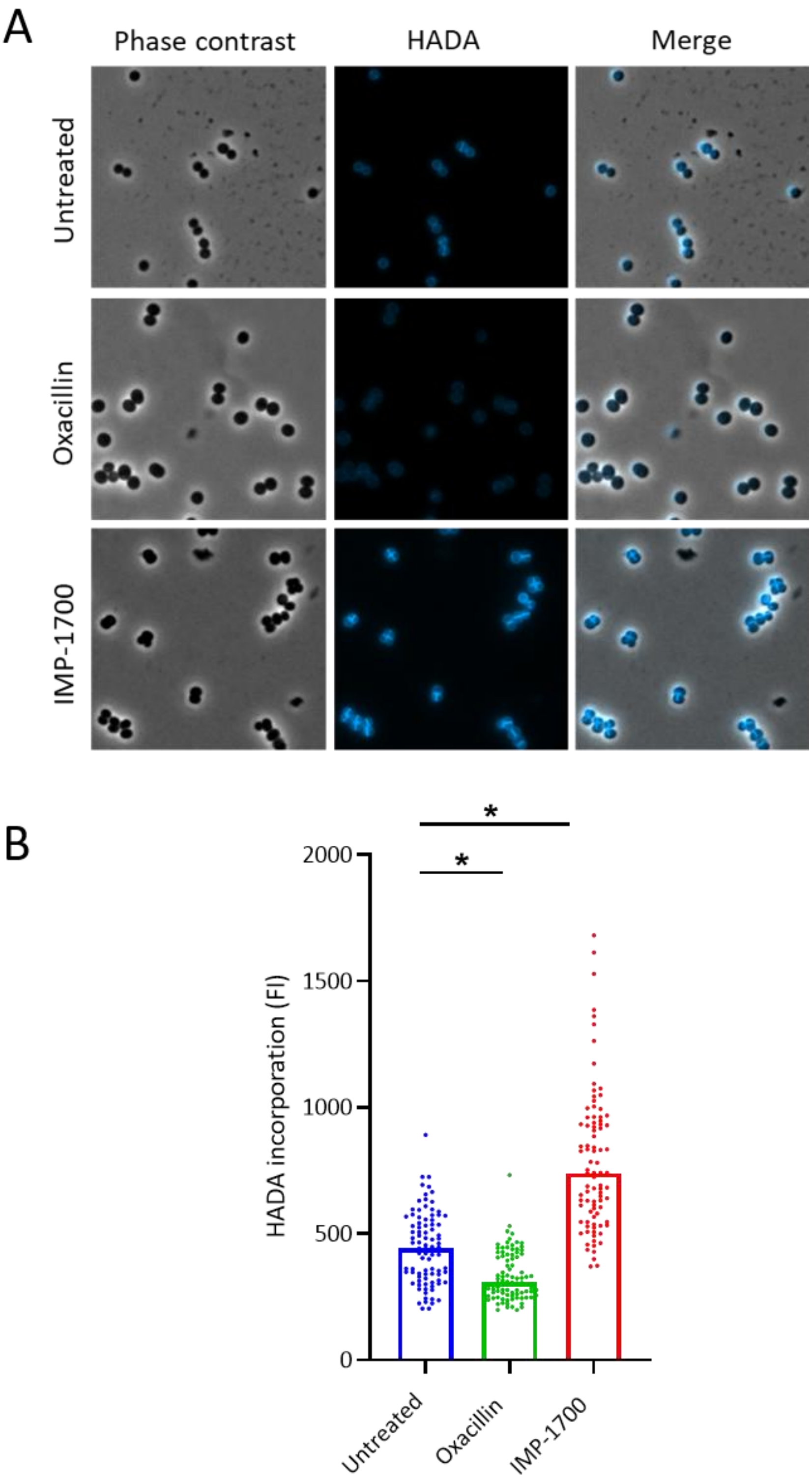

**Supplementary Figure S3. IMP-1700 increases septal wall biosynthesis in MRSA. (A)** Incorporation of HADA into the cell wall of *S. aureus* JE2 WT was observed by phase contrast and fluorescence microscopy. Cultures were incubated with 100 µM HADA and either left untreated or exposed to IMP-1700 or oxacillin (2 X MIC) to inhibit peptidoglycan synthesis, for 2 h. Scale bar, 2 µm. **(B)** HADA fluorescence intensity of the septum was quantified using FIJI software. Data represents 30 cells per biological replicate (90 cells total) with the median of each replicate shown. Analyses were performed using a one-way ANOVA with Dunnett's post-hoc test (*p < 0.01 relative to untreated).

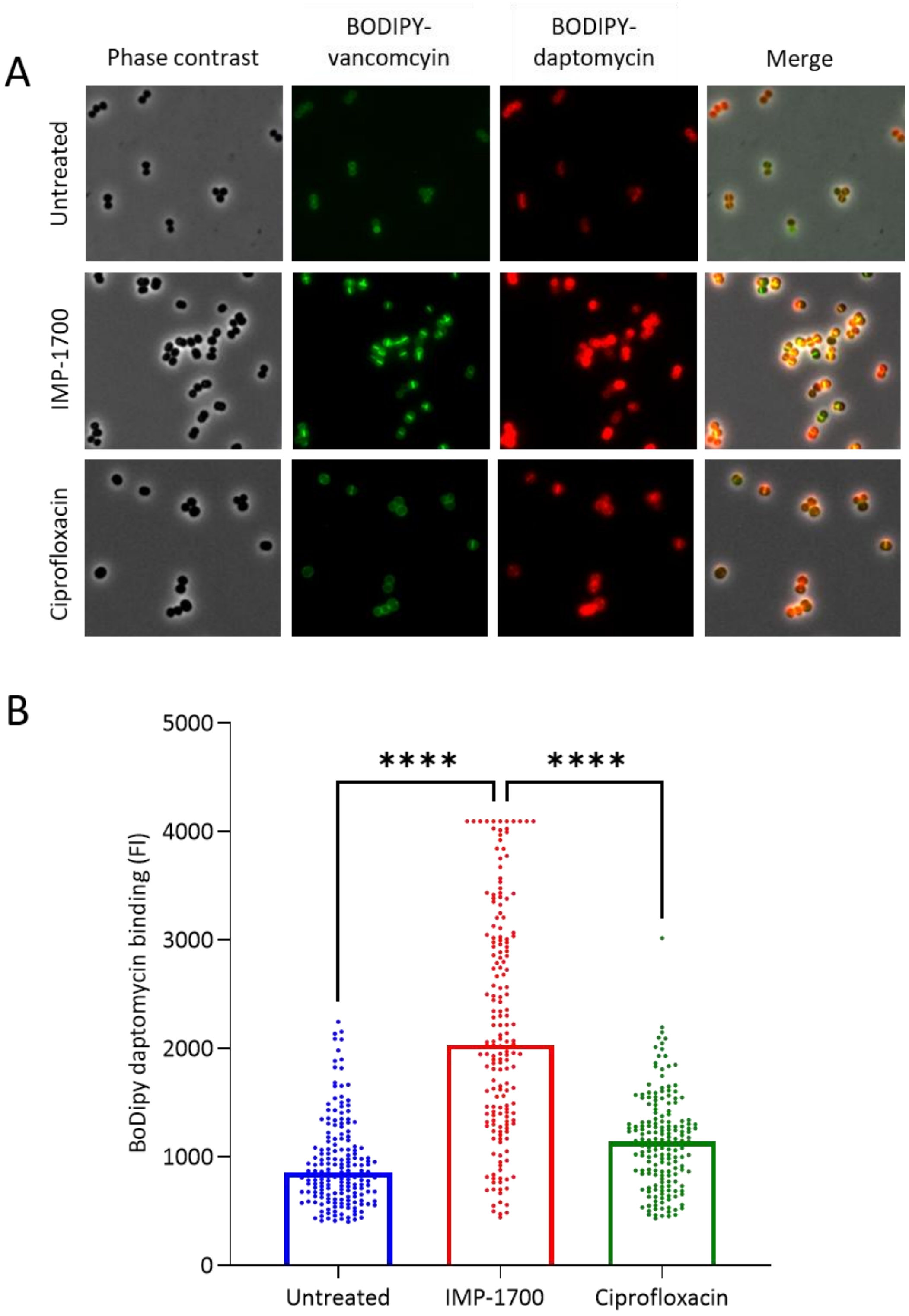

**Supplementary Figure S4. IMP-1700 increases daptomycin binding in *S. aureus* JE2. (A)** *S. aureus* JE2 was incubated with 2X MIC of IMP-1700, ciprofloxacin or neither for 2h before staining with BODIPY-vancomycin and BODIPY-daptomycin. **(B)** BODIPY-daptomycin binding of cells in **(A)**. The fluorescence intensity was derived from a line drawn perpendicular to the septum of cells stained with BODIPY-daptomycin and data represents the mean ± standard deviation of 20 cells per replicate (n=3, 60 cells in total).

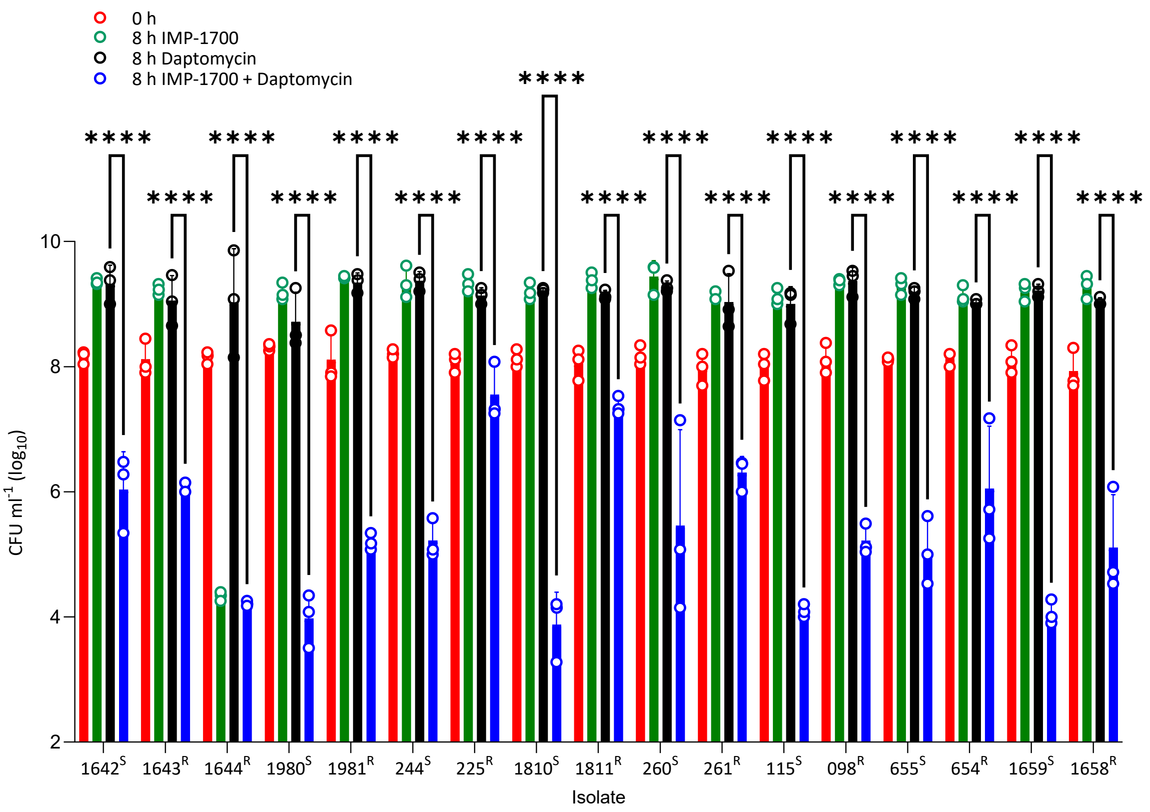

**Supplementary Figure S5. IMP-1700 potentiates the bactericidal activity of daptomycin against clinical daptomycin-susceptible and non-susceptible *S. aureus* isolates.** Isolates were exposed to 2X MIC IMP-1700 either alone or in combination with daptomycin (5 µg ml$^{-1}$) for 8 h. Most isolates grew in the presence of IMP-1700 or daptomycin alone, despite >MIC concentrations due to the size of the starting inoculum, which is 200-fold greater than that used in MIC assays. Data shown are the mean ± standard deviation of three independent experiments analysed by a two-way ANOVA with Dunnet's post-hoc test (****P < 0.0001 relative to daptomycin alone).
